# Quantitative Economon Model of Transactions for Drugs and Other Commodities

**DOI:** 10.1101/2025.07.08.663082

**Authors:** Jeremy M. Haynes, Joel Correa da Rosa, Eduardo R. Butelman, Steven R. Hursh, S. Stevens Negus

## Abstract

**Background:** An economon is a dyadic economic unit of two participants who exchange mutually reinforcing commodities (e.g., addictive drugs for money). A human economon often consists of a buyer providing money to a supplier, while the supplier reciprocally provides some commodity to the buyer. Here, we develop the Quantitative Economon Model (QEM) by reviewing basic principles of behavioral economics and translating those principles into a quantitative model characterizing transactional behavior between buyers and suppliers that would be applicable to transactions involving drugs specifically, as well as other commodities. According to the model, transactions between a buyer and supplier depend on their respective economic demand for the commodities exchanged. Additionally, the model assumes that demand for commodities fluctuates across sequential transactions due to satiation and deprivation associated with consumption of those commodities, of particular relevance to addictive drugs.

**Methods:** We used a computational approach to simulate data from the QEM according to four conditions representing parametric manipulations of high or low levels of deprivation crossed with high or low levels of satiation (i.e., a 2×2 matrix of conditions).

**Results:** Simulations revealed patterns of transactional behavior that may be characteristic of a range of commodities. In particular, when deprivation-effects were high and satiation-effects were low, simulated transaction rates were characteristic of the high consumption rates observed with addictive drugs. In other conditions, transaction rates varied along patterns characteristic of consumption for other commodities like food. In sum, the QEM provides a framework for modeling, investigating, and predicting patterns of human transactional behavior.

## 1 Introduction

Addictions (substance use disorders; SUD) are diseases of consummatory behavior characterized by both (1) excessive consumption of one commodity (i.e., a specific drug) to the point of causing harm and (2) neglect of other commodities and associated activities that could promote health (e.g., healthy food, prosocial behaviors & activities of daily living; American Psychiatric Association, 2013). However, the emergence of addiction depends not only on behavior by the consumer or “buyer” of the addictive commodity, but also on actions by the supplier of that commodity (Hoffer et al., 2009; Hursh, 1991; Hursh & Schwartz, 2023; Negus, 2024). This broader contextual focus (Acuff et al., 2024) thus includes commercial and social networks in the development of SUD in specific environments (Boivin, 2013; National Institute on Drug Abuse, 2024; Sharma et al., 2017). Moreover, drug addiction is a uniquely human pathology because, to our knowledge, humans are the only species capable not only of responding to the rewarding and reinforcing effects of addictive substances (in common with several other species; Ahmed et al., 2002; Negus, 2006; Zernig et al., 2007; Zhang et al., 2015), *but also of producing and selling those substances*. We have suggested that the interactive, dyadic, and mutually reinforcing relationship (Reid & Mukhopadhyay, 2021) between the buyer and supplier of addictive as well as non-addictive commodities might best be considered together in a transactional unit labeled an “economon” (plural: economa; Negus, 2024). As illustrated in Fig. 1A, the simplest form of an economon consists of two participants who have an opportunity to exchange mutually reinforcing commodities. In the case of an addiction-focused economon, the buyer obtains the addictive substance (the reinforcer for the buyer) from a supplier, and the supplier in turn obtains money or some other commodity (the reinforcer for the supplier) from the buyer (Negus, 2024) .

**Fig. 1.**
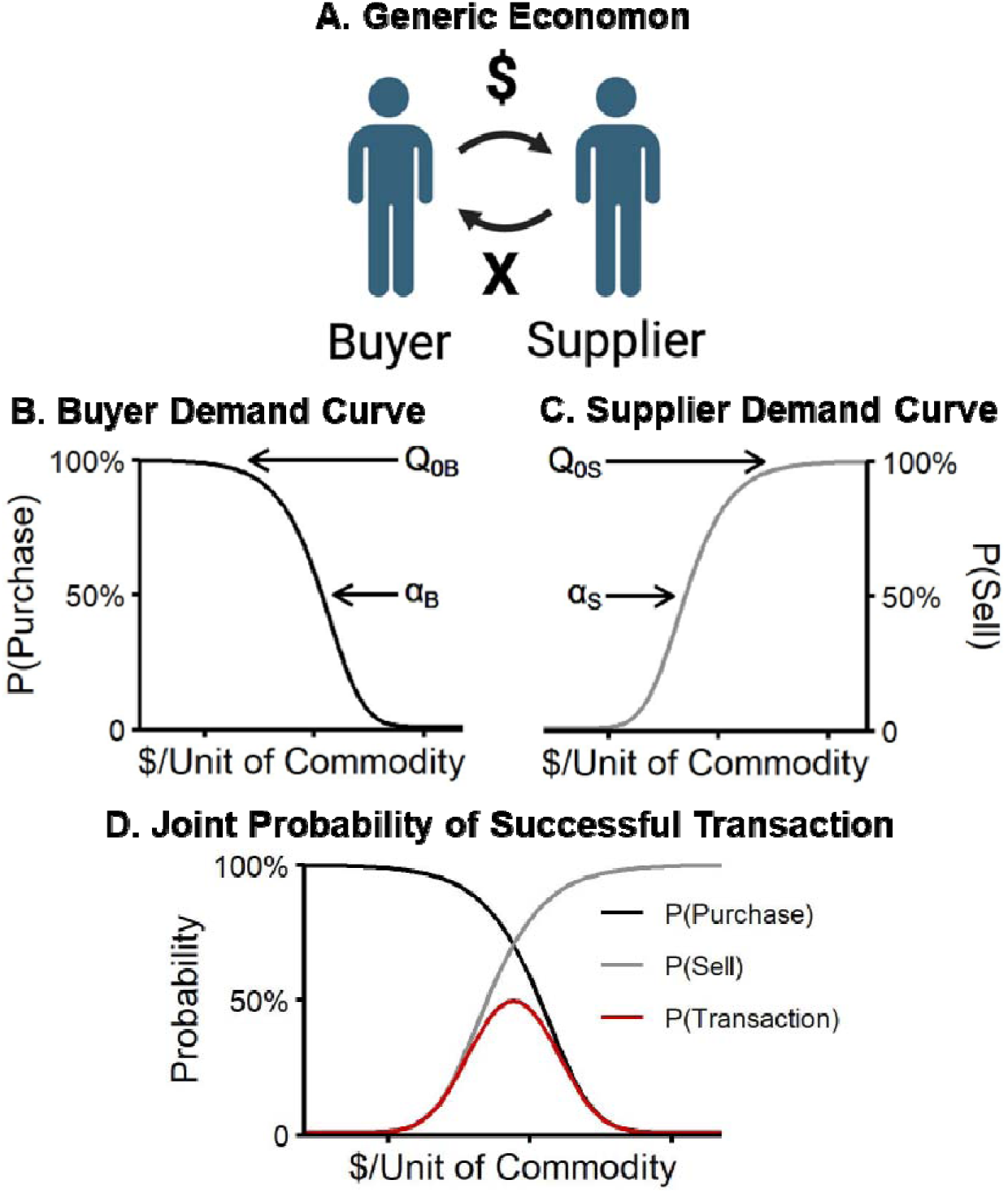
Conceptual Model of Economon Framework. *Note.* A: Dyadic relationship between buyer and supplier exchanging money and commodity “X” (e.g., drug), labeled an economon. B: Buyer demand curve according to Equation 1. C: Supplier demand curve according to Equation 2. D: Overlapping buyer and supplier demand curves with transaction probability (red datapath) determined by the joint probability of the buyer purchasing the commodity and the supplier selling the commodity according to Equation 3.

Although this conceptual description of an economon can be helpful in exploring factors related to the emergence of addiction (Negus, 2024), a quantitative implementation of the model would foster more precise definitions of the factors that govern economon operation, enable simulation of the impact of those factors on buyer and supplier behaviors over time, and guide empirical efforts to measure, predict, and ultimately intervene on those factors. Therefore, here we extended the economon framework by developing the Quantitative Economon Model (QEM) to quantitatively characterize transactional behavior involving drug commodities as well as a range of transactional behaviors that could also include non-drug commodities, as will be illustrated below. Accordingly, the major goal of this manuscript is to describe the basic features of the QEM and to initiate the use of this dyadic agent-based model to simulate human economic behavior (Bonabeau, 2002; Hoffer et al., 2009). To do this, we first developed the quantitative framework of the model, assuming that such a model should account for both buyer *and* supplier behavior individually and jointly, as well as how transactions between the buyer and supplier change across time in response to whether the buyer and supplier exchange the commodities.

Next, we used a computational (i.e., simulation-based) approach to generate predictions from the model based on specific variations of model parameters. Based on predictions from the model, we describe the emergent properties of the model’s operation according to those variations in parameter values.

## 2 Methods

### 2.1 Parameters of the Model

As described above, the Quantitative Economon Model (QEM) accounts for the behavior of individual buyers and suppliers across sequential encounters. As the starting point of the model, we review and use the well-established behavioral economic concept of a demand curve

(Hursh & Silberberg, 2008). As shown in Fig. 1B, the demand curve for the buyer of a commodity (e.g., an addictive drug) relates that commodity’s unit price (i.e., $/unit of commodity) on the *x* axis to the demand for the commodity on the *y* axis. Demand typically refers to the amount of a commodity that a buyer will purchase at a given price (Hursh et al., 2013); however, in Fig. 1B, and for all following demand functions, we represent demand on the *y* axis in terms of the hypothetical probability (Jacobs & Bickel, 1999) of buying a commodity at a given price (Hayashi et al., 2022). In general, increases in price lead to decreases in demand and the associated likelihood of buying the commodity, and the resulting curve is characterized by two parameters: *Q_0_*, which defines a theoretical rate of maximal demand of the commodity when price = 0 (i.e., intensity of demand, represented here as 100% likelihood of buying), and α, which defines the elasticity of demand for the commodity (i.e., the rate at which demand decreases with increases in price, or “price sensitivity”; Hursh et al., 2013). Together, *Q_0_* and α describe the shape of a demand curve such that demand increases with increases in *Q_0_* and with decreases in α, or demand decreases with decreases in *Q_0_* and with increases in α. Fig. 1B was generated with a simplified version of Hursh and Silberberg’s exponential demand function for the buyer (Hursh & Silberberg, 2008; see also Hursh & Schwartz, 2023; Winger et al., 2007, 2006):

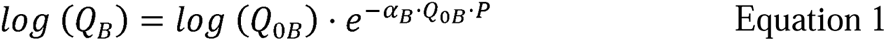

where *Q_B_* represents buyer demand at unit price *P* (i.e., $ / unit of the commodity), and *Q_0B_* and α*_B_* are as described above with the subscript *B* specifically indicating the buyer. Thus, Equation 1 characterizes how increases in unit price are associated with decreases in demand and the associated probability of purchasing the commodity.

The QEM builds on the demand-curve concept in three novel ways. First, the QEM stipulates that supplier behavior in an economon can also be represented by a demand curve. That is, supplier demand for money (the reinforcer to the supplier) as a function of the commodity being supplied (i.e., the cost to the supplier) can also be expressed by a demand curve. For the supplier, unit price refers to “units of commodity / $” (i.e., cost / reinforcer), which is the reciprocal of unit price for the buyer. However, Fig. 1 C shows that we can represent the demand curve for the supplier as a function of unit price for the buyer, so that supplier demand can be plotted on the same *x* axis continuum as the buyer demand curve. To do this, we simply take unit price for the supplier, units of commodity / $, and take the reciprocal to get $ / unit of commodity. Thus, for both the buyer and supplier, prices arrayed on the *x* axis of the demand curve are expressed as a ratio (e.g. $ / mg of a drug such as heroin or fentanyl), with the numerator and denominator of this ratio describing the commodities being exchanged. As shown in Fig. 1B, the buyer trades the numerator commodity (money) to receive the denominator commodity (e.g., heroin), and buyer price increases from left to right along the *x* axis.

Conversely, as shown in Fig. 1C, the supplier trades the denominator commodity to receive the numerator commodity (e.g. heroin for dollars), and supplier price increases from *right to left* along the *x* axis. Mathematically, the demand curve for the supplier, displayed in Fig. 1C, is represented as the following:

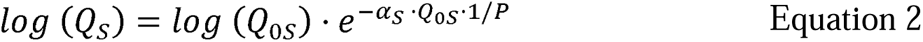

such that *Q_S_* represents supplier demand at unit price 1/*P*, and *Q_0S_* and α*_S_* are the supplier *Q_0_*and α parameters, respectively, with the subscript *S* specifically indicating the supplier. The resulting supplier demand curve is shown in Fig. 1C, and its general shape (demand declining as supplier price rises from *right to left*) mirrors the buyer demand curve (demand declining as buyer price rises from left to right).

These buyer and supplier demand curves are shown separately in Figs. 1B and 1C for initial clarity, but they can also be plotted together as shown in Fig. 1D, and this leads to the second way in which the QEM builds on the demand-curve concept. Specifically, the QEM makes the working assumption that the success of a transaction depends on *both* the buyer agreeing to purchase the commodity *and* the supplier agreeing to sell the commodity at a given price, as postulated in standard economic models (Jenkin, 1870). Stated mathematically, the probability of a successful transaction is based on the joint probability of the buyer and supplier demand curves at a given price:

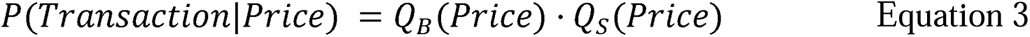

where *Q_B_(Price)* and *Q_S_(Price)* are derived from Equations 1 and 2, respectively. At very low prices, the buyer has a high probability of purchase, but the supplier has a negligible probability of sale. Conversely, at very high prices, the supplier has a high probability of sale, but the buyer has a negligible probability of purchase. As shown by the red curve in Fig. 1D, the probability of a successful transaction therefore peaks at intermediate prices where buyer and supplier demand curves overlap and both buyer and supplier are likely to participate in the transaction.

Lastly, the QEM assumes that demand curves are not static within individual buyers or suppliers but instead can fluctuate across sequential encounters due to “satiation” or “deprivation” for the exchanged commodities in response to transaction outcomes and the associated consumption or lack of consumption of those commodities. Here, we operationalize consumption and lack of consumption between encounters as the success or failure of a previous transaction, respectively. Thus, successful transactions would result in the delivery of the respective commodities to each economon participant and produce some level of satiation to decrease demand during subsequent encounters. For example, “satiation” from food intake can be conceptualized as a composite variable with different bio-behavioral components (Amin & Mercer, 2016; Tremblay & Bellisle, 2015) that reduces demand for food in a time-dependent manner (e.g., based on time proximity to consumption). Based on archival data (Greenwald & Hursh, 2006; Hursh et al., 1989), deprivation and satiation primarily affect elasticity of demand (i.e., α) with little effect on the intensity of demand (i.e., *Q_0_*; see supplement for further discussion). Thus, we represent satiation-related changes in demand as occurring due to increases in elasticity:

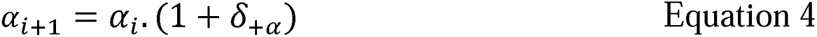

where a*_i_* is elasticity before encounter *i*, and δ*_+_*_D_ (constrained: 0 ≤ δ*_+_*_α_) is a parameter that characterizes proportional *increases* in elasticity. For example, a δ*_+_*_α_ = .75 would correspond to an increase in elasticity by 75%, resulting in decreases in demand on subsequent encounters.

Conversely, failed transactions would result in some level of deprivation that increases demand via decreases in elasticity during subsequent encounters. Using a food commodity as an example again, deprivation from lack of food intake may increase demand for food as the duration of deprivation increases (Amin & Mercer, 2016; Casanova et al., 2019). As with satiation-related changes in elasticity, we represent deprivation-related changes as decreases in elasticity:

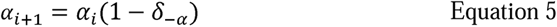

where a*_i_* is elasticity before encounter *i*, and δ*_-_*_α_ (constrained: 0 ≤ δ*_-_*_α_ ≤ 1) is a parameter that characterizes proportional *decreases* in elasticity. For example, a δ*_-_*_α_ = .75 would correspond with a decrease in elasticity by 75%, resulting in increases in demand on subsequent encounters. Thus, these two alternative outcomes (successful vs. failed transaction) are incorporated into the model by including two additional parameters for each participant that characterize changes in elasticity of demand for the buyer and the supplier. Table 1 summarizes the parameters for the model.

**Table 1.** Parameters for Quantitative Economon Model.

| Parameter | Description |
| --- | --- |
| $Q_{0B}$<br>$Q_{0S}$ | Theoretical maximal probability that the buyer ($B$) will purchase the commodity when price (\$ / unit of commodity) is 0, and that the supplier ($S$) will sell the commodity when the price is infinite. |
| $\alpha_B$<br>$\alpha_S$ | Elasticity of demand for buyer ($B$) and supplier ($S$). That is, the rate at which increases in price produce decreases in probability of engaging in the transaction. |
| $\delta_{+aB}$<br>$\delta_{+aS}$ | Proportional *increase* in elasticity for the buyer ($B$) and supplier ($S$) after a successful transaction, described as satiation. |
| $\delta_{-aB}$<br>$\delta_{-aS}$ | Proportional *decrease* in elasticity for the buyer ($B$) and supplier ($S$) after a failed transaction, described as deprivation. |

### 2.2 Operation of the Model

The model operates by arranging a series of sequential buyer/supplier encounters, with each encounter resulting in a transaction outcome that updates buyer and supplier demand curves for the next encounter. Thus, each encounter provides the opportunity for a transaction, with the outcome of that transaction determined by buyer and supplier demand curves and the resulting joint probability of a transaction as shown by the red datapath in Fig. 1D (Equation 3). Operation of the model is depicted in Fig. 2. At the beginning of an encounter, participants enter with their respective demand curves. Based on the likelihood of purchasing/selling the commodity (derived from Equations 1 & 2), Bernoulli distributions determine buyer and supplier engagement in a transaction at a given price. When both participants agree to a transaction, then the commodities are exchanged; otherwise, the commodities are not exchanged. In the simulation below, we implement this process using the probabilities of purchasing (Equation 1) and selling (Equation 2) the commodity in two independent Bernoulli distributions, but this process could also be implemented using the joint probability of purchasing and selling the commodity (Equation 3) in a single Bernoulli distribution, with identical results. After an encounter, the buyer and supplier demand curves are updated to reflect satiation (after a successful transaction) or deprivation (after a failed transaction) with Equations 4 and 5. The effects of deprivation and satiation on buyer and supplier demand across encounters are illustrated in Fig. 3. A successful transaction produces satiation-related increases in buyer and supplier α, moving the demand curves farther apart, reducing overlap in the demand curves, and thus also reducing the probability of a successful transaction on the next encounter. Conversely, a failed transaction produces deprivation-related decreases in buyer and supplier α, moving demand curves toward each other, producing more overlap, and thus increasing the probability of a successful transaction on the next encounter. Over a series of sequential encounters, buyer and supplier demand curves will migrate toward and away from each other in response to failed or successful transactions, respectively.

**Fig. 2.**
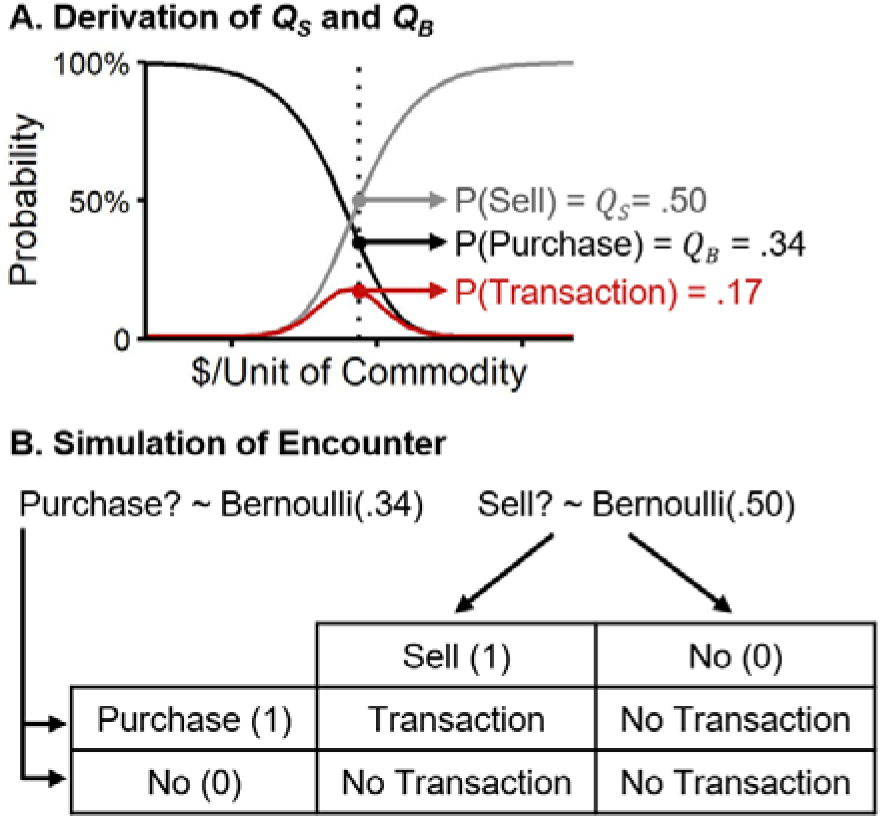
Diagram of Simulation. *Note.* Framework for economon operation during the simulation. A: Upon an encounter between a buyer and supplier, probabilities of purchasing and selling the commodity were calculated from the buyer and supplier demand curves (i.e., Equations 1-2), respectively, at a set unit price (dotted vertical line). Price was constant across encounters and set at the price corresponding to ½ of the supplier *Q_0S_*. B: Using the purchase and sell probabilities calculated in A, Bernoulli distributions were used to determine whether the buyer and seller agree (1) or do not agree (0) to buy and sell the commodity at a given price, respectively. A transaction is successful when both agents agree; otherwise, a transaction fails.

**Fig. 3.**
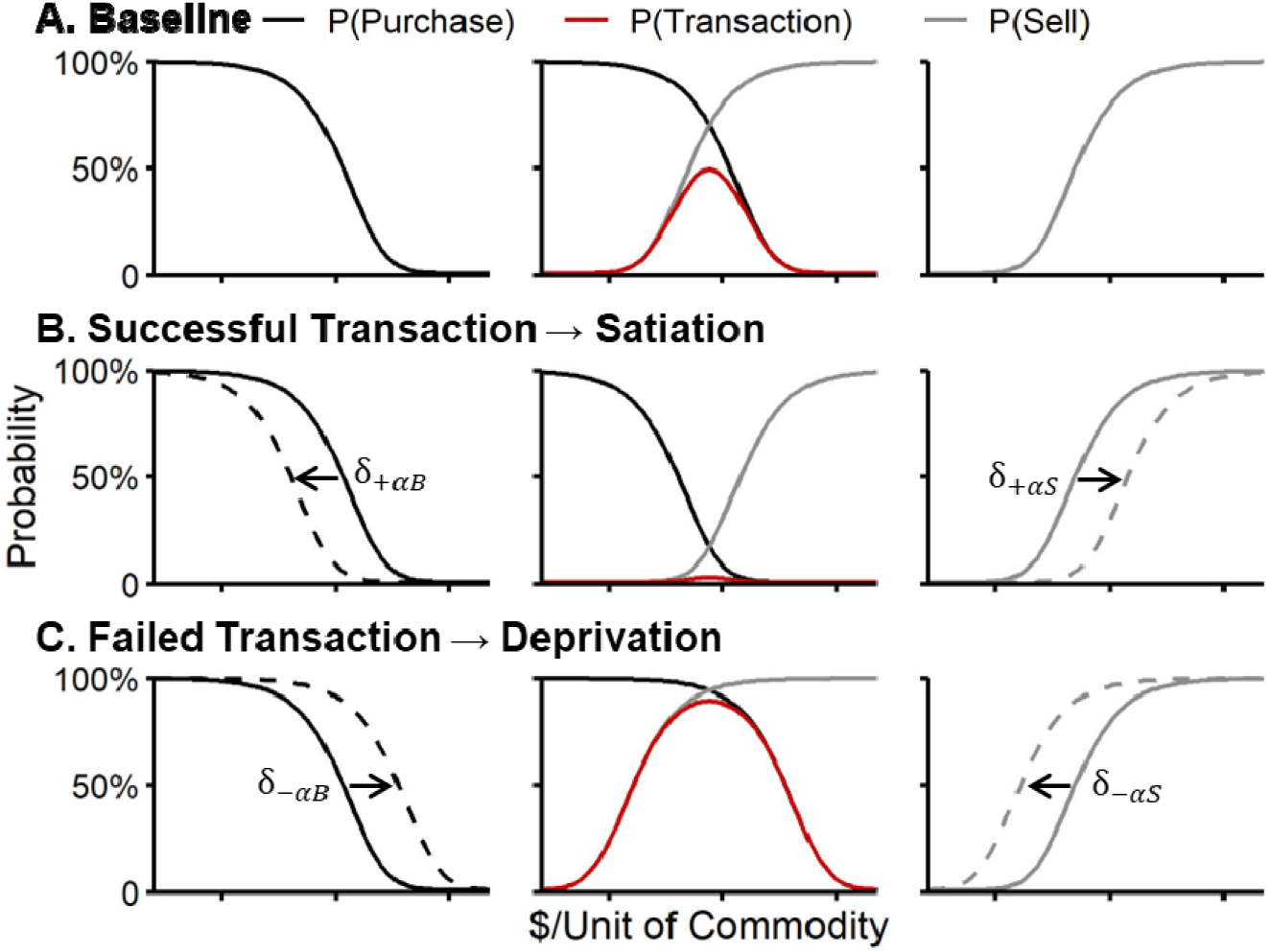
Changes in Demand Following Successful & Failed Transactions. *Note.* Probability of purchasing and selling a commodity (e.g., drug) by a buyer (left column) and supplier (right column), respectively, as a function of commodity price (i.e., $ / unit of commodity). The middle column overlays buyer and supplier demand from adjacent columns, with the joint probability depicted by the red datapath, representing the probability of a successful transaction for that encounter. A: Baseline demand at the start of an encounter, used to calculate the joint probability of a successful transaction. B: Demand following a successful transaction (dashed datapath) in which demand decreases, from baseline (solid datapath), for both the buyer and supplier via satiation-induced increases in elasticity, determined by δ*_+_*_α_. C: Demand following a failed transaction (dashed datapath) in which demand increases, from baseline (solid datapath), for both the buyer and supplier via deprivation-induced decreases in elasticity, determined by δ*_-_*_α_.

Given the iterative nature of how elasticities change across encounters based on failed and successful transactions, we used a computational (i.e., simulation-based) approach to characterize the emergent properties of the model. For simplicity, this initial application of the model limits parametric manipulation in three ways. First, we only manipulated buyer parameters (δ*_+_*_α_*_B_* & δ*_-_*_α_*_B_*) across conditions, while holding supplier parameters constant (i.e., δ*_+_*_α_*_S_* = 0 & δ*_-_*_α_*_S_* = 0) across conditions. Second, parametric manipulations were not examined across the entire range of prices, but instead at a single price, set arbitrarily at ½ of the supplier *Q_0S_*. Finally, we selected specific “high” and “low” values for the buyer satiation and deprivation parameters, shown in Table 2, rather than examining a full range of parameter values. After simulating data, we summarized transaction patterns across sequential encounters to identify the emergent properties of each condition.

**Table 2.** Group-Level Deprivation and Satiation Parameters for Buyers in Simulation.

| Condition | Deprivation | Satiation | Example of commodity purchased |
| --- | --- | --- | --- |
| Low Dep / High Sat | $\delta_{-aB} = .075$ | $\delta_{+aB} = 4$ | Durable good (e.g. car) |
| Low Dep / Low Sat | $\delta_{-aB} = .075$ | $\delta_{+aB} = 1$ | Durable luxury good (e.g., jewelry) |
| High Dep / High Sat | $\delta_{-aB} = .500$ | $\delta_{+aB} = 4$ | Transient essential good (e.g. food) |
| High Dep / Low Sat | $\delta_{-aB} = .500$ | $\delta_{+aB} = 1$ | Transient inessential good (e.g. fentanyl for an individual with OUD) |

*Note.* Seller parameters were held constant in simulation. OUD: opioid use disorder; Dep: deprivation; Sat: satiation.

### 2.3 Computational Approach for the QEM

To illustrate QEM operation, we simulated transactions between buyers and suppliers across sequential encounters under four selected conditions for the satiation and deprivation parameters (described below). Fig. 2 shows the simulation process for individual encounters. In an encounter, to determine whether a transaction succeeded or failed, we used Equations 1-2 to calculate the probability of the buyer purchasing the commodity (*Q_B_*) and the supplier selling the commodity (*Q_S_*), respectively (Fig. 2A). The probabilities were used to randomly generate each agent’s decision on engaging in a transaction, according to a Bernoulli distribution:

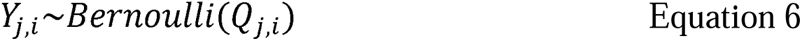

where *Y_j,i_* E {0,1} refers to whether the buyer (*j* = *B*) or supplier (*j* = *S*) agree to a transaction on the *i*-th encounter (Fig. 2B). The transaction in the *i*-th encounter is successful if both agents agree, *Y_B,i_* = 1 and *Y_S,i_*= 1, otherwise the transaction is unsuccessful (i.e., no transaction). This corresponds to an outcome from a bivariate Bernoulli distribution (Marshall & Olkin, 1985).

As described above, we held the supplier’s demand constant (i.e., no deprivation or satiation effects over the sequence of encounters); however, buyer demand varied from encounter-to-encounter according to δ*_+_*_α_*_B_* and δ*_-_*_α_*_B_*. We selected two levels of δ*_+_*_α_*_B_* (“high” vs. “low” satiation effects) crossed with two levels of δ*_-_*_α_*_B_*, (“high” vs. “low” deprivation effects), resulting in four conditions. Table 2 shows the parameter values for each of these conditions. Within each condition, we simulated 700 sequential encounters between a buyer and supplier for 100 hypothetical economa (i.e. 100 different buyer-supplier dyads for each condition). To introduce variability across economa, we added random error to the δ*_+_*_α_*_B_* and δ*_-_*_α_*_B_* parameters according to normal distributions with means of 0 and standard deviations of 1.0 and 0.1, respectively. Although rare, if an individual δ*_+_*_α_*_B_* and δ*_-_*_α_*_B_* fell outside of the parameter’s bounds (noted above), we constrained the individual δ*_+_*_α_*_B_* and δ*_-_*_α_*_B_* parameters to the values displayed in Table 2. For economa in each condition, we removed the first 200 encounters between the buyer and supplier as “preconditioning” to avoid inclusion of encounters that were not representative of the stable transaction patterns that emerged for each condition (cf. “burn-in” for a Markov Chain Monte Carlo algorithm; Kruschke, 2010). After these “burn-in” samples were removed, the remaining 500 were visually checked for ‘stability’ (i.e., no increasing/decreasing trends in group-level purchase probabilities). Here, we only present and summarize results for the first 100 samples after preconditioning (i.e., samples 201 to 300) for visual simplicity. These samples are representative of the full dataset, which is shown in the supplemental file. Because supplier demand was held constant, we focus our visual analysis on purchase probabilities from buyers across each condition.

Code is freely available on GitHub (model: https://github.com/correadarosaj/Economon.git) and OSF (simulation & supplement: https://osf.io/uc6yr/?view_only=71a3117ab1194e1dafb9b5c432333a7e).

## 3 Results

Overall, we observed distinct patterns of transactions across the four conditions of the simulation, reflecting how the deprivation and satiation parameters affected elasticity of demand for the buyer from encounter to encounter. To illustrate, Fig. 4 displays group level (i.e., mean) purchase probabilities across encounters for each condition. Summary statistics of the group-level data are presented in Table 3. In addition, we highlight person-level data from a representative buyer within each panel of Fig. 4, and we display demand curves for a subset of transaction opportunities from those buyers in Fig. 5. These buyers were selected because their δ*_+_*_α_ and δ*_-_*_α_ values were closest to the parameter values presented in Table 2. The full simulation is displayed in the supplement (Fig. S1), which also includes an animation illustrating demand across encounters during the simulation, for representative conditions (Fig. S2).

**Fig. 4.**
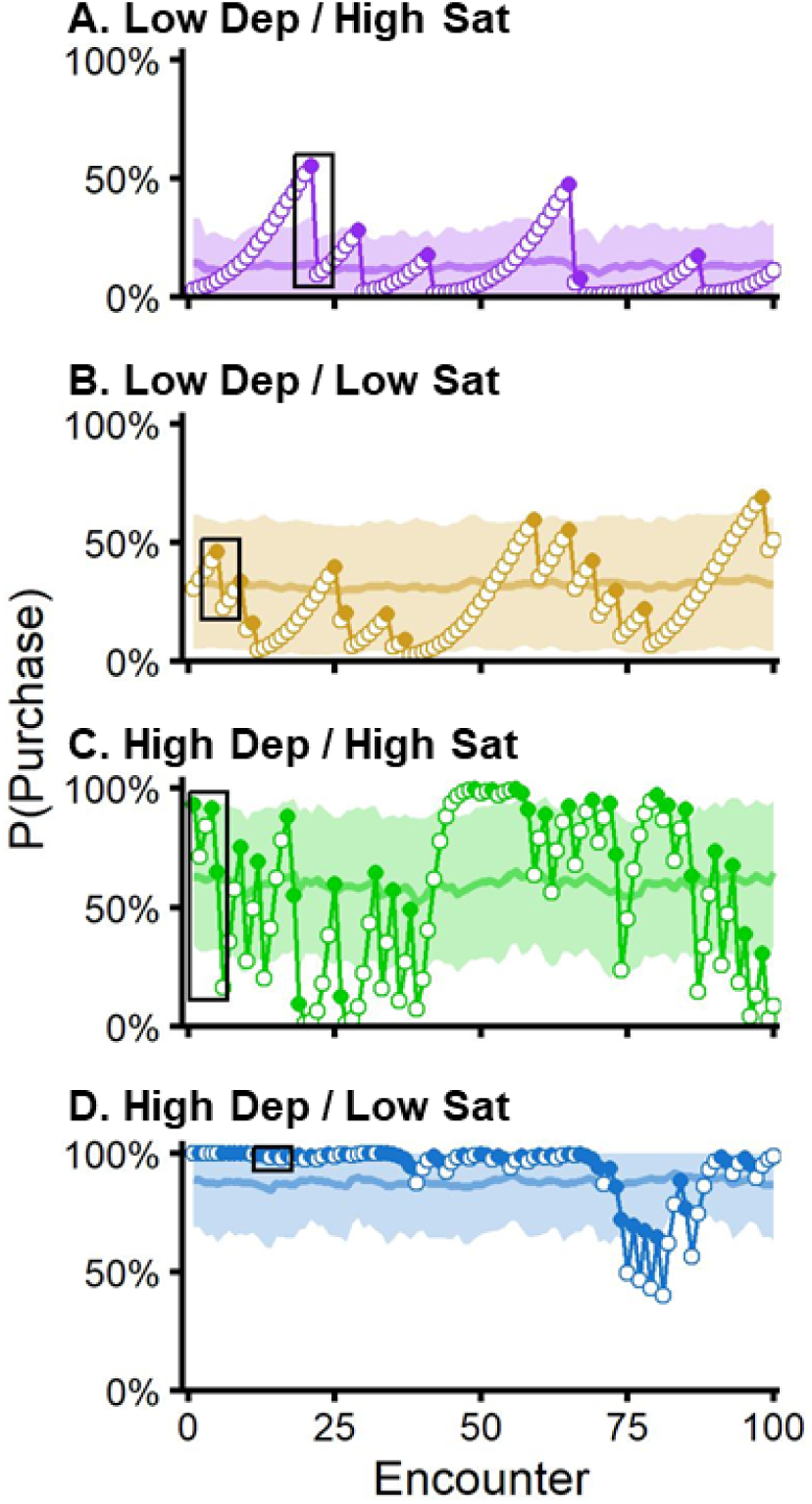
Purchase Probabilities from Simulation. *Note.* Probability of the simulated buyers purchasing a commodity as a function of encounters with a supplier. Purchase probabilities changed according to failed and successful transactions. Failed transactions caused increases in demand for the commodity (due to deprivation); successful transactions caused decreases in demand for the commodity (due to satiation). Light datapaths represent group level (i.e., mean) purchase probabilities from 100 buyers across 100 encounters with suppliers. Shaded areas represent ±SD around the means. Darker datapaths represent person level purchase probabilities from one representative buyer within each condition. The inset in each panel indicates encounters presented in Fig. 5.

**Fig. 5.**
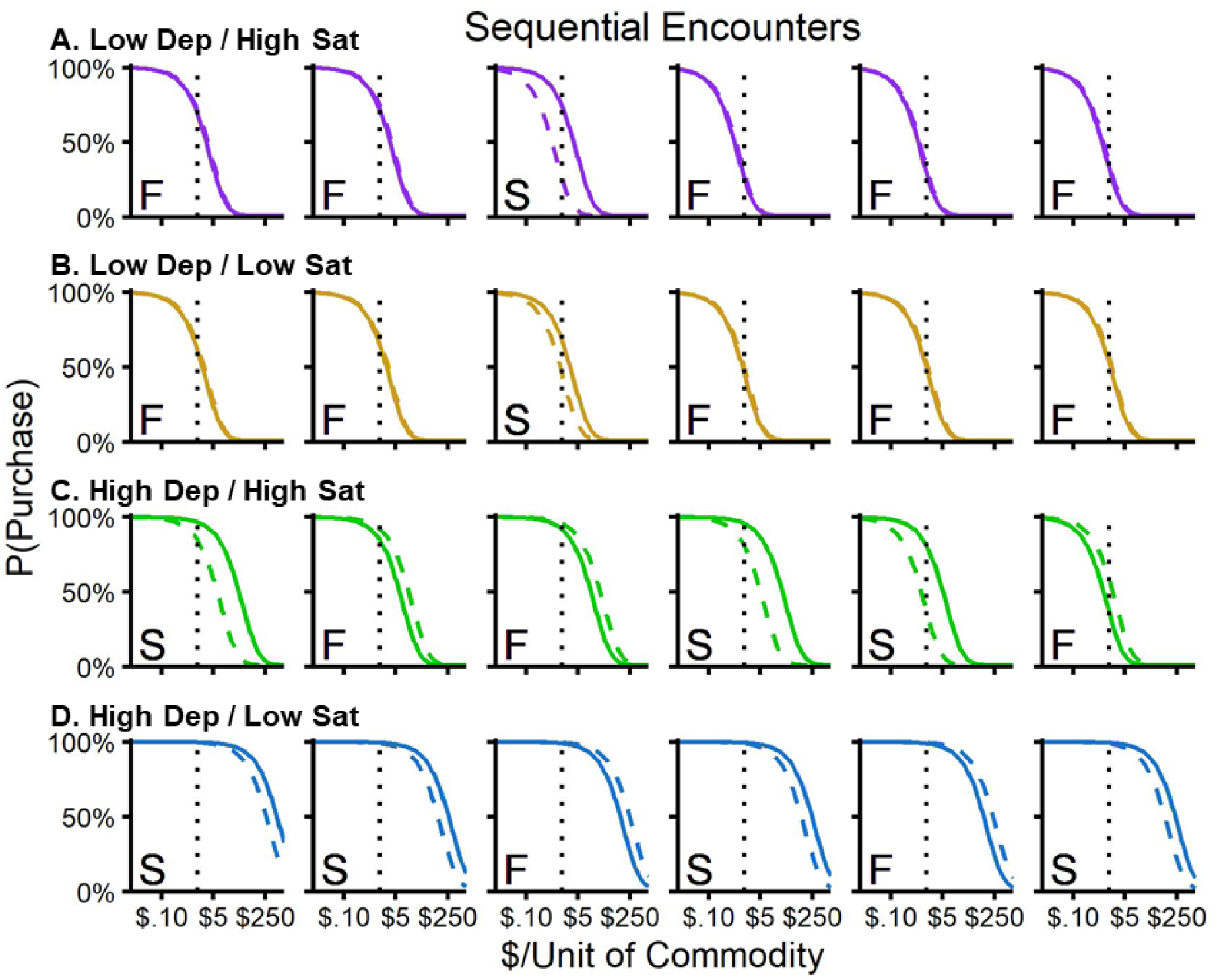
Demand Curves from Simulation. *Note.* Changes in demand for six selected sequential encounters for a representative buyer from each condition (rows), for illustration. Solid datapaths correspond to demand before the encounter and dashed datapaths correspond to demand after the encounter (i.e., after deprivation- or satiation-related changes in demand). Price of the commodity during the encounter is represented by the dotted vertical line, which also represents the price at ½ *Q_0S_*. Transaction outcomes were determined at this price for all simulations. F: failed transaction; S: successful transaction.

**Table 3.** Summary Statistics from Simulation.

| Condition | $\alpha_B$ | $Q_B$ | Transaction Rate | Latency |
| --- | --- | --- | --- | --- |
| Low Dep / High Sat | .00940<br>(.05167) | .13<br>(.16) | 6.73<br>(4.10) | 13.81<br>(14.36) |
| Low Dep / Low Sat | .00195<br>(.00271) | .32<br>(.28) | 15.91<br>(11.11) | 5.27<br>(7.17) |
| High Dep / High Sat | .00086<br>(.00176) | .60<br>(.32) | 30.19<br>(6.23) | 2.32<br>(2.28) |
| High Dep / Low Sat | .00016<br>(.00045) | .88<br>(.21) | 43.69<br>(8.00) | 1.30<br>(1.68) |

*Note.* Means (standard deviations) calculated across simulations for each condition. Transaction Rate: number of successful transactions across the 100 encounters displayed in Fig. 4; Latency: number of trials between successful transactions for the 100 encounters displayed in Fig. 4.

As Fig. 4A and Table 3 show, the Low Dep / High Sat condition had the lowest purchase probability on average and generated the lowest transaction rate. At the person level, the likelihood of purchasing the commodity was cyclical, such that purchase probabilities steadily increased across failed transactions (open symbols) until a transaction occurred (filled symbols), at which time the purchase probability dropped drastically (see also Fig. 5A Column 3). For the Low Dep / Low Sat condition (Fig. 4B), the average purchase probability and transaction rate (see Table 3) was higher than for the Low Dep / High Sat condition (above). Person-level purchase probabilities for the representative buyer from the Low Dep / Low Sat condition were also cyclical. As Fig. 5B shows, demand changed little from encounter to encounter in this condition. The High Dep / High Sat condition (Fig. 4C) had a higher purchase probability and transaction rate than the previous two conditions, as well as greater variability in demand from encounter to encounter. As Fig. 5C shows, demand during the High Dep / High Sat condition changed more drastically across successful and failed transactions. Finally, the High Dep / Low Sat condition (Fig. 4D) resulted in the highest average purchase probability and transaction rate compared to each of the other conditions but also showed relatively little variability in demand from encounter to encounter (see Fig. 5D). Within the High Dep / Low Sat condition, we observed that, occasionally, a long series of sequential successful transactions resulted in transient decreases in demand (e.g., in the individual datapath in Fig. 4D). The full simulation, displayed in the supplement (Fig. S1), shows this pattern approximately once every 150 encounters, thus representing a sporadic characteristic of this condition. Combined, results from the simulation show that varying levels of the deprivation and satiation parameters in the QEM result in distinct patterns of transactions across encounters between economa participants (i.e., buyers & suppliers of a commodity). Next, we discuss the implications of the model and future directions.

## 4 Discussion

This is the first quantitative implementation of the economon model (Negus, 2024), which focuses on sequential encounters in a dyad composed of a buyer and a supplier. This model has the potential to model transaction patterns for any exchanged commodity in future implementations, based on an agent-based framework (Ahearne et al., 2022). To our knowledge, this is one of the few models designed to examine person-level buying and selling interactions in a scalable manner, amenable to examination of parameters that can be directly measured based on expanding understanding of, for example, substance use disorder (SUD; Acuff et al., 2024; Bobashev et al., 2020; McLellan et al., 2022). In prior SUD research, agent-based models have primarily, but not exclusively (Hoffer et al., 2009), focused on macro-level processes (e.g., group-level interactions between buyers & suppliers or healthcare systems; Hursh, 1991), or longer-term interactions (Bobashev et al., 2020; Di Clemente & Pietronero, 2012; Lim et al., 2022; Mollick & Kober, 2020; Tatara et al., 2024; Zhong et al., 2023). The novel features and flexibility of the Quantitative Economon Model (QEM) are designed to allow for hypothesis generation and testing of properties related to transactional behavior in general, and in the trajectories of addiction and other economic pathologies in individuals and populations in particular (Epstein, 2008).

### 4.1 Emergent Phenotypes from the QEM simulation

We implemented the QEM based on principles from behavioral economics that characterize demand for a commodity according to two parameters: (1) the intensity of demand (*Q_0_*) and (2) elasticity of demand (α; Hursh and Silberberg, 2008). For the QEM, we extended these traditional behavioral economic principles by assuming that supplier behavior can also be expressed by a demand curve in which suppliers pay with a commodity to receive money, but we express their economic behavior in terms of unit price for the buyer (i.e., $ / unit of commodity) so that both of these demand curves can be displayed on the same *x* axes. In addition, we make the working assumption that the likelihood of a “successful” (i.e., completed) transaction between a buyer and supplier depends on both agents agreeing to buy and sell the commodity at a given price. Finally, the model assumes that demand elasticity, and thus demand itself, of both buyers and suppliers changes dynamically depending on whether a prior encounter resulted in a successful or failed transaction. In the model, we characterized these changes in elasticity by parameters related to satiation (δ*_+_*_α_) and deprivation (δ*_-_*_α_; Chen et al., 2023). In this initial implementation, we used a computational approach to simulate patterns of transactional behavior and determined emergent properties of the model according to four conditions that differed with respect to two selected levels of satiation (“high” & “low”) crossed with two selected levels (“high” & “low”) of deprivation for the buyer while holding the supplier’s demand constant (see Table S1 in supplement). Below, we summarize and discuss the results of the simulations for each condition.

#### 4.1.1 Low Deprivation / High Satiation

The condition with low deprivation and high satiation was characterized by failed transactions resulting in small decreases in elasticity, and successful transactions resulting in large increases in elasticity. At the group level, this resulted in the fewest number of successful transactions because buyer demand was relatively low (i.e., low probability of purchase). At the person level, this condition resulted in orderly patterns of transactions characterized by demand steadily increasing across encounters due to sequential failed transactions, until a successful transaction occurred, at which time demand decreased dramatically to low levels. These patterns of transactions were cyclical across encounters, resulting in more failed than successful transactions. Commodities associated with these patterns of economic behavior could include relatively essential commodities with long lifespans, such as a car (Cho et al., 2024).

#### 4.1.2 Low Deprivation / Low Satiation

The condition with low deprivation and low satiation was also characterized by failed transactions resulting in small decreases in elasticity, but with successful transactions resulting in small increases in elasticity. At the group level, this condition was associated with more successful transactions than that of the Low Deprivation / High Satiation condition (above) but was still associated with fewer successful than failed transactions on average. At the person level, this condition also resulted in orderly patterns of transactions. The cyclical patterns of transactions were similar to the Low Deprivation / High Satiation condition, but because successful transactions produced smaller satiation-related decreases in demand, successful transactions were more likely in general and in closer proximity with one another (see encounters between 60-75 in Fig. 4B). Commodities associated with these patterns of economic behavior could include inessential but desirable (i.e., luxury) commodities with long lifespans, such as articles of jewelry.

#### 4.1.3 High Deprivation / High Satiation

The condition with high deprivation and high satiation was characterized by failed transactions resulting in large decreases in elasticity and successful transactions resulting in large increases in elasticity. Combined, these changes in elasticity produced a high degree of variability in demand (i.e., probability of purchase) for the commodity. The higher variability in demand was not only observed at the group level, but also at the person level where demand varied widely from encounter to encounter (see Fig. 4C). Despite the variability, this condition had a high rate of transactions, with over a 50% likelihood of purchasing the commodity on average at the group level. In addition, this condition did not show the cyclical changes in demand as the previous two conditions showed (i.e., the conditions with low deprivation parameters). Commodities associated with these patterns of economic behavior could include essential commodities such as food (de-Magistris & Gracia, 2017; Otterbring et al., 2024), which have short lifespans and generally homeostatically-maintained satiation and deprivation mechanisms (Amin & Mercer, 2016; Tremblay & Bellisle, 2015).

#### 4.1.4 High Deprivation / Low Satiation

Finally, the condition with high deprivation and low satiation was also characterized by failed transactions resulting in large decreases in elasticity, but with successful transactions resulting in relatively small increases in elasticity. At the group level, this condition was associated with the most transactions because demand was high across encounters in this condition. Likewise, demand was also high across encounters at the person level (near 100% likelihood of purchasing the commodity); however, occasionally, a series of sequential successful transactions resulted in demand falling to surprisingly low levels (<50%) until recovering to nearly a 100% likelihood of purchasing the commodity. This was not a sampling issue (i.e., failure of the Markov chain Monte Carlo algorithm reaching steady state) but instead appeared to be an emergent property of this combination of parameters (see Fig. S1).

Commodities associated with these patterns of economic behavior could include inessential but desirable commodities with short lifespans. For example, in persons chronically exposed to fentanyl or heroin, such as those diagnosed with opioid use disorder, lack of consumption may result in rapid and severe deprivation due to dependence, and consumption may not result in a high level of satiation due to tolerance (Pergolizzi et al., 2020).

### 4.2 Opportunities for QEM Elaboration

The four conditions described above represent simple case scenarios tentatively associated with representative commodities; however, additional parametric manipulations are possible within the model framework which will be necessary to more adequately characterize buyer-supplier behaviors in real-world economa with empirical data for addictive substances (e.g., Table S1 in supplement). Below, we discuss four possible elaborations of the QEM.

#### 4.2.1 Transaction Outcome-Effects on Supplier Elasticity

First, for the simplified simulations in the present study, only buyer elasticity (α*_B_*) was allowed to fluctuate after successful/failed transactions, while supplier elasticity (α_S_) was held constant. Assuming that many suppliers receive money, which due to its fungibility is less susceptible to deprivation- and satiation-related changes in demand compared with other commodities (Charlton and Fantino, 2008), we consider this a reasonable starting point.

However, in a more complete model, supplier elasticity would also be affected by transaction outcomes. For example, in barter economies where goods and services are exchanged for other goods and services, rather than for money, the magnitude of transaction outcome-effects on supplier elasticity would depend on the commodity being received by the supplier (e.g., sex for providing a drug; Rash et al., 2016).

#### 4.2.2 Transaction Outcome-Effects on *Q_0_*

Second, demand curves are defined not only by the α parameter, but also by *Q_0_*(peak demand at the lowest price). For the present simulations, *Q_0_*was set arbitrarily at 100% for both participants and was unaffected by transaction outcomes, similar to *normalized demand* (Hursh and Winger, 1995). In a more complete model, baseline *Q_0_* could lie at any point along the 0-100% probability continuum depending on the commodity and the preference of the participant for that commodity. For example, the peak purchase probability might be low for a commodity that a buyer does not prefer, even if that commodity is essentially free (i.e., with little-to-no cost). Moreover, this *Q_0_* could also be sensitive to transaction outcomes because consumption of a commodity may “carry over” (Baucells & Sarin, 2007) to influence the motor and/or cognitive competence of the consumer to engage in further consumption (van Steenbergen et al., 2019).

For example, consumption of an opioid drug like fentanyl or heroin may produce transient sedation that would reduce motor competence in a time-dependent manner, possibly reflected by a lower *Q_0_*.

#### 4.2.3 Consumption-Induced Changes in Deprivation-/Satiation-Effects

Third, in the present simulation, transaction outcomes produced constant changes in elasticity regardless of prior consumption levels. That is, the degree to which satiation (δ*_+_*_α_) and deprivation (δ*_-_*_α_) affected elasticity of demand did not fluctuate across encounters. However, repeated consumption of commodities like addictive drugs can produce changes in buyer biology and neurocognitive effects that accrue to affect the degree of deprivation and satiation over time (e.g., dependence & withdrawal; Kearns et al., 2011; Koob, 2019; Koob & Kreek, 2007). For example, repeated consumption of opioid drugs may produce both physical dependence leading to gradual increases in the degree of deprivation (i.e., increases in withdrawal or dysphoria) across buyer-supplier encounters (Hall et al., 2024; Koob, 2019) which could be characterized by increases in δ*_-_*_α_. Likewise, gradual decreases in the degree of satiation across repeated consumption of opioid drugs (i.e., development of tolerance & escalation; Zernig et al., 2007) could be characterized by decreases in δ*_+_*_α_. Thus, changes in the degree of deprivation and/or satiation could be modeled as changes in δ*_+_*_α_ and δ*_-_*_α_, respectively, based on prior consumption (Chen et al., 2023).

#### 4.2.4 Moving from Isolated Economa to Networks of Economa

Lastly, the present study considered economa operating in isolation (i.e., independent buyer-seller dyads). However, economa in the real world exist in networks (i.e., interact with one another; Kranton & Minehart, 2001) such that the operation of one economon can influence the operation of other economa in its network. Importantly, the degree to which economa interact, and the nature of these interactions, within a network may serve as indicators of health-related problems. In a simple example, a buyer may have access to two suppliers offering two different commodities (e.g. an addictive drug vs. food), and the extent to which one economon produces more successful transactions relative to another (e.g., drug economon over food economon) can be indicative of overall health and diagnoses (e.g., for substance use disorders; Hasin et al., 2013). Also, the presence of substitutes by different suppliers (e.g., fentanyl vs. heroin, which have different pharmacological potency; Comer et al., 2008) should affect the operation of economa, both in terms of buyer behavior (i.e., allocation to one supplier over another) and supplier behavior (i.e., competition; Hursh, 1991). Addressing the four issues described above would allow for a more extensive and nuanced application of the QEM to characterize trajectories of consumption for maladaptive commodities such as addictive drugs.

### 4.3 Opportunities for Empirical Research

Ultimately, our goal in developing the Quantitative Economon Model (QEM) is to guide experimental design and data collection for future empirical studies that examine transactional behavior between two agents. In its current form, the QEM highlights four buyer and supplier variables determining transactional behavior: (1) intensity of demand (*Q_0_*), (2) elasticity of demand (α), (3) satiation-related increases in demand elasticity (δ*_+_*_α_), and (3) deprivation-related decreases in demand elasticity (δ*_-_*_α_). Accordingly, a major goal for economon-based research going forward will involve the laboratory- or field-based measurement of these variables for both buyers and suppliers, and currently available research tools can be used to accomplish this goal. For example, hypothetical purchase tasks (HPTs) are questionnaires that ask human subjects to predict their level of consumption for a given commodity across a range of prices, and the resulting data can be used to generate demand curves with associated *Q_0_* and α parameters (Jacobs & Bickel, 1999; Mackillop et al., 2008; Murphy & MacKillop, 2006; Strickland & Lacy, 2020; see also Gilroy et al., 2025). HPTs have been used to assess demand of both drug and non-drug reinforcers (Epstein et al., 2018; Strickland et al., 2020; Strickland & Stoops, 2017), and prevailing evidence suggests good correspondence between hypothetical demand assessed with HPTs and actual demand assessed in tasks that involve actual purchases and consumption (González-Roz et al., 2019; Kaplan et al., 2018; Martínez-Loredo et al., 2021; Strickland & Lacy, 2020). Because HPTs can be completed rapidly (∼10 min), they could be used to assess *Q_0_* and α parameters before and after sequential transaction opportunities to measure changes in those parameters after successful vs. failed transactions. Similarly, ecological momentary assessments (EMAs) use mobile devices, wearable sensors, and other technology to enable real-time data collection from subjects in natural settings (Preston et al., 2009; Preston & Epstein, 2011). Therefore, EMAs could be used to obtain time-dependent “real world” data on self-rated demand and the magnitude of satiation- or deprivation-related changes in demand after drug consumption. These empirically determined data could then be used to refine estimates of QEM parameters (*Q_0_*, α, δ*_+_*_α_ & δ*_-_*_α_) for drugs and other commodities in future simulations and strengthen the real-world relevance. Importantly, understanding this transactional behavior between buyers and suppliers could have important implications for targeting interventions aimed to improve human health (e.g., reduce substance abuse; National Institute on Drug Abuse, 2024).

### 4.5 Summary

In sum, the Quantitative Economon Model (QEM) provides a computational representation of the interaction between two agents engaging in mutual reinforcement, based on principles derived from behavioral economics. For many economa, a buyer will exchange money to receive a commodity (e.g., drug) and a supplier will exchange the commodity to receive money. According to the model, the probability that such an exchange occurs depends on the economic demand of both agents for their respective reinforcers (i.e., the commodity for the buyer & money for the supplier). The model further assumes that demand is not constant across time, but rather, depends on whether previous encounters between a buyer and supplier result in a successful or failed transaction, causing satiation- or deprivation-related changes in demand.

Using this computational implementation, we characterized the emergent properties of the model according to different levels of deprivation and satiation for a buyer engaging with a supplier in an economon. Overall, the degree to which deprivation and satiation affected demand was a primary determinant of the frequency and pattern of successful transactions. Although further research is necessary to empirically examine specific aspects of the QEM, the model provides a foundation for understanding economa and their implications for human health problems such as addiction.

## References

Acuff, S.F., Strickland, J.C., Smith, K., Field, M., 2024. Heterogeneity in choice models of addiction: the role of context. Psychopharmacology (Berl.) 241, 1757–1769. 10.1007/s00213-024-06646-1

Ahearne, M., Atefi, Y., Lam, S.K., Pourmasoudi, M., 2022. The future of buyer–seller interactions: a conceptual framework and research agenda. Journal of the Academy of Marketing Science 50, 22–45. 10.1007/s11747-021-00803-0

Ahmed, S.H., Kenny, P.J., Koob, G.F., Markou, A., 2002. Neurobiological evidence for hedonic allostasis associated with escalating cocaine use. Nat. Neurosci. 5, 625–626. 10.1038/nn872

American Psychiatric Association, D., 2013. . academia.edu.

Amin, T., Mercer, J.G., 2016. Hunger and satiety mechanisms and their potential exploitation in the regulation of food intake. Curr. Obes. Rep. 5, 106–112. 10.1007/s13679-015-0184-5

Baucells, M., Sarin, R.K., 2007. Satiation in discounted utility. Oper. Res. 55, 170–181. 10.1287/opre.1060.0322

Bobashev, G.V., Hoffer, L.D., Lamy, F.R., 2020. Agent-based modeling to delineate opioid and other drug use epidemics, in: Complex Systems and Population Health. Oxford University PressNew York, pp. 201–C14.P78. 10.1093/oso/9780190880743.003.0014

Boivin, R., 2013. Risks, prices, and positions: A social network analysis of illegal drug trafficking in the world-economy. Int. J. Drug Policy. 10.1016/j.drugpo.2013.12.004

Bonabeau, E., 2002. Agent-based modeling: methods and techniques for simulating human systems. Proc. Natl. Acad. Sci. U. S. A. 99 Suppl 3, 7280–7287. 10.1073/pnas.082080899

Casanova, N., Finlayson, G., Blundell, J.E., Hopkins, M., 2019. Biopsychology of human appetite — understanding the excitatory and inhibitory mechanisms of homeostatic control. Curr. Opin. Physiol. 12, 33–38. 10.1016/j.cophys.2019.06.007

Charlton, S.R., Fantino, E., 2008. Commodity specific rates of temporal discounting: does metabolic function underlie differences in rates of discounting? Behav. Processes 77, 334–342. 10.1016/j.beproc.2007.08.002

Chen, W., He, Y., Bansal, S., 2023. Customized Dynamic Pricing when customers develop a habit or satiation. Oper. Res. 71, 2158–2174. 10.1287/opre.2022.2412

Cho, C., Matsumoto, B., Smith, D., 2024. A consumption measure for automobiles. Mon. Labor Rev. 10.21916/mlr.2024.3

Comer, S.D., Sullivan, M.A., Whittington, R.A., Vosburg, S.K., Kowalczyk, W.J., 2008. Abuse liability of prescription opioids compared to heroin in morphine-maintained heroin abusers. Neuropsychopharmacology 33, 1179–1191. 10.1038/sj.npp.1301479

de-Magistris, T., Gracia, A., 2017. Does hunger matter in consumer purchase decisions? An empirical investigation of processed food products. Food Qual. Prefer. 55, 1–5. 10.1016/j.foodqual.2016.08.002

Di Clemente, R., Pietronero, L., 2012. Statistical Agent Based Modelization of the Phenomenon of Drug Abuse. Sci. Rep. 2, 1–8. 10.1038/srep00532

Epstein, J.M., 2008. Why model? Journal of artificial societies and social simulation 11, 12.

Epstein, L.H., Paluch, R.A., Carr, K.A., Temple, J.L., Bickel, W.K., MacKillop, J., 2018. Reinforcing value and hypothetical behavioral economic demand for food and their relation to BMI. Eat. Behav. 29, 120–127. 10.1016/j.eatbeh.2018.03.008

Gilroy, S.P., Rzeszutek, M.J., Koffarnus, M.N., Reed, D.D., Hursh, S.R., 2025. Adaptive purchase tasks in the operant demand framework. Exp. Clin. Psychopharmacol. 10.1037/pha0000757

González-Roz, A., Jackson, J., Murphy, C., Rohsenow, D.J., MacKillop, J., 2019. Behavioral economic tobacco demand in relation to cigarette consumption and nicotine dependence: a meta-analysis of cross-sectional relationships. Addiction 114, 1926–1940. 10.1111/add.14736

Greenwald, M.K., Hursh, S.R., 2006. Behavioral economic analysis of opioid consumption in heroin-dependent individuals: effects of unit price and pre-session drug supply. Drug Alcohol Depend. 85, 35–48. 10.1016/j.drugalcdep.2006.03.007

Hall, O.T., Gunawan, T., Teater, J., Bryan, C., Gorka, S., Ramchandani, V.A., 2024. Withdrawal interference scale: a novel measure of withdrawal-related life disruption in opioid use disorder and alcohol use disorder. Am. J. Drug Alcohol Abuse 1–13. 10.1080/00952990.2024.2350057

Hasin, D.S., O’Brien, C.P., Auriacombe, M., Borges, G., Bucholz, K., Budney, A., Compton, W.M., Crowley, T., Ling, W., Petry, N.M., Schuckit, M., Grant, B.F., 2013. DSM-5 criteria for substance use disorders: recommendations and rationale. Am. J. Psychiatry 170, 834– 851. 10.1176/appi.ajp.2013.12060782

Hayashi, Y., Fisher, N.M., Hantula, D.A., Furman, L., Washio, Y., 2022. A behavioral economic demand analysis of mothers’ decision to exclusively breastfeed in the workplace. J. Exp. Anal. Behav. 118, 132–147. 10.1002/jeab.772

Hoffer, L.D., Bobashev, G., Morris, R.J., 2009. Researching a local heroin market as a complex adaptive system. Am. J. Community Psychol. 44, 273–286. 10.1007/s10464-009-9268-2

Hursh, S.R., 1991. Behavioral economics of drug self-administration and drug abuse policy. J. Exp. Anal. Behav. 56, 377–393. 10.1901/jeab.1991.56-377

Hursh, S.R., Madden, G.J., Spiga, R., DeLeon, I.G., Francisco, M.T., 2013. The translational utility of behavioral economics: The experimental analysis of consumption and choice, in: APA Handbook of Behavior Analysis, Vol. 2: Translating Principles into Practice. American Psychological Association, Washington, pp. 191–224. 10.1037/13938-008

Hursh, S.R., Raslear, T.G., Bauman, R., Black, H., 1989. The quantitative analysis of economic behavior with laboratory animals, in: Understanding Economic Behaviour. Springer Netherlands, Dordrecht, pp. 393–407. 10.1007/978-94-009-2470-3_22

Hursh, S.R., Schwartz, L.P., 2023. A general model of demand and discounting. Psychol. Addict. Behav. 37, 37–56. 10.1037/adb0000848

Hursh, S.R., Silberberg, A., 2008. Economic demand and essential value. Psychol. Rev. 115, 186–198. 10.1037/0033-295X.115.1.186

Hursh, S.R., Winger, G., 1995. Normalized demand for drugs and other reinforcers. J. Exp. Anal. Behav. 64, 373–384.

Jacobs, E.A., Bickel, W.K., 1999. Modeling drug consumption in the clinic using simulation procedures: demand for heroin and cigarettes in opioid-dependent outpatients. Exp. Clin. Psychopharmacol. 7, 412–426. 10.1037//1064-1297.7.4.412

Jenkin, F., 1870. The graphic representation of the laws of supply and demand, and their application to labour, in: Grant, A. (Ed.), Recess Studies. Edmonston & Douglas, Edinburgh, pp. 151–170.

Kaplan, B.A., Foster, R.N.S., Reed, D.D., Amlung, M., Murphy, J.G., MacKillop, J., 2018. Understanding alcohol motivation using the alcohol purchase task: A methodological systematic review. Drug Alcohol Depend. 191, 117–140. 10.1016/j.drugalcdep.2018.06.029

Kearns, D.N., Gomez-Serrano, M.A., Tunstall, B.J., 2011. A review of preclinical research demonstrating that drug and non-drug reinforcers differentially affect behavior. Curr. Drug Abuse Rev. 4, 261–269. 10.2174/1874473711104040261

Koob, G.F., 2019. Neurobiology of Opioid Addiction: Opponent Process, Hyperkatifeia, and Negative Reinforcement. Biol. Psychiatry. 10.1016/j.biopsych.2019.05.023

Koob, G., Kreek, M.J., 2007. Stress, dysregulation of drug reward pathways, and the transition to drug dependence. Am. J. Psychiatry 164, 1149–1159. 10.1176/appi.ajp.2007.05030503

Kranton, R.E., Minehart, D.F., 2001. A theory of buyer-seller networks. Am. Econ. Rev. 91, 485–508. 10.1257/aer.91.3.485

Kruschke, J., 2010. Doing Bayesian Data Analysis: A tutorial with R and BUGS. Europe’s Journal of Psychology 7, 778–779. 10.5964/EJOP.V7I4.163

Lim, T.Y., Stringfellow, E.J., Stafford, C.A., DiGennaro, C., Homer, J.B., Wakeland, W., Eggers, S.L., Kazemi, R., Glos, L., Ewing, E.G., Bannister, C.B., Humphreys, K., Throckmorton, D.C., Jalali, M.S., 2022. Modeling the evolution of the US opioid crisis for national policy development. Proc. Natl. Acad. Sci. U. S. A. 119, e2115714119. 10.1073/pnas.2115714119

Mackillop, J., Murphy, J.G., Ray, L.A., Eisenberg, D.T., Lisman, S.A., Lum, J.K., Wilson, D.S., 2008. Further validation of a cigarette purchase task for assessing the relative reinforcing efficacy of nicotine in college smokers. Experimental and clinical psychopharmacology 16.

Marshall, A.W., Olkin, I., 1985. A family of bivariate distributions generated by the bivariate Bernoulli distribution. J. Am. Stat. Assoc. 80, 332–338. 10.1080/01621459.1985.10478116

Martínez-Loredo, V., González-Roz, A., Secades-Villa, R., Fernández-Hermida, J.R., MacKillop, J., 2021. Concurrent validity of the Alcohol Purchase Task for measuring the reinforcing efficacy of alcohol: an updated systematic review and meta-analysis. Addiction 116, 2635–2650. 10.1111/add.15379

McElreath, R., 2020. Statistical rethinking, 2nd ed. CRC Press, London, England. 10.1201/9780429029608

McLellan, A.T., Koob, G.F., Volkow, N.D., 2022. Preaddiction-A Missing Concept for Treating Substance Use Disorders. JAMA Psychiatry 79, 749–751. 10.1001/jamapsychiatry.2022.1652

Mollick, J.A., Kober, H., 2020. Computational models of drug use and addiction: A review. J. Abnorm. Psychol. 129, 544–555. 10.1037/abn0000503

Murphy, J.G., MacKillop, J., 2006. Relative reinforcing efficacy of alcohol among college student drinkers. Exp. Clin. Psychopharmacol. 14, 219–227. 10.1037/1064-1297.14.2.219

National Institute on Drug Abuse, 2024. Commercial interests contribute to drug use and addiction [WWW Document].

National Institute on Drug Abuse. URL https://nida.nih.gov/about-nida/noras-blog/2024/09/commercial-interests-contribute-to-drug-use-addiction (accessed 9.28.24).

Negus, S.S., 2024. An economon model of drug addiction. Psychopharmacology . 10.1007/s00213-024-06535-7

Negus, S.S., 2006. Choice between heroin and food in nondependent and heroin-dependent rhesus monkeys: effects of naloxone, buprenorphine, and methadone. J. Pharmacol. Exp. Ther. 317, 711–723. 10.1124/jpet.105.095380

Otterbring, T., Folwarczny, M., Gasiorowska, A., 2024. The impact of hunger on indulgent food choices is moderated by healthy eating concerns. Front. Nutr. 11, 1377120. 10.3389/fnut.2024.1377120

Pergolizzi, J.V., Jr, Raffa, R.B., Rosenblatt, M.H., 2020. Opioid withdrawal symptoms, a consequence of chronic opioid use and opioid use disorder: Current understanding and approaches to management. J. Clin. Pharm. Ther. 45, 892–903. 10.1111/jcpt.13114

Preston, K.L., Epstein, D.H., 2011. Stress in the daily lives of cocaine and heroin users: relationship to mood, craving, relapse triggers, and cocaine use. Psychopharmacology (Berl.) 218, 29–37. 10.1007/s00213-011-2183-x

Preston, K.L., Vahabzadeh, M., Schmittner, J., Lin, J.-L., Gorelick, D.A., Epstein, D.H., 2009. Cocaine craving and use during daily life. Psychopharmacology (Berl.) 207, 291–301. 10.1007/s00213-009-1655-8

Rash, C.J., Burki, M., Montezuma-Rusca, J.M., Petry, N.M., 2016. A retrospective and prospective analysis of trading sex for drugs or money in women substance abuse treatment patients. Drug Alcohol Depend. 162, 182–189. 10.1016/j.drugalcdep.2016.03.006

Reid, C., Mukhopadhyay, S., 2021. Mutual reinforcement learning with heterogenous agents, in: 2021 IEEE International Conference on Smart Computing (SMARTCOMP). Presented at the 2021 IEEE International Conference on Smart Computing (SMARTCOMP), IEEE. 10.1109/smartcomp52413.2021.00081

Sharma, A., Sinha, K., Vandenberg, B., 2017. Pricing as a means of controlling alcohol consumption. Br. Med. Bull. 123, 149–158. 10.1093/bmb/ldx020

Strickland, J.C., Campbell, E.M., Lile, J.A., Stoops, W.W., 2020. Utilizing the commodity purchase task to evaluate behavioral economic demand for illicit substances: a review and meta-analysis. Addiction 115, 393–406. 10.1111/add.14792

Strickland, J.C., Lacy, R.T., 2020. Behavioral economic demand as a unifying language for addiction science: Promoting collaboration and integration of animal and human models. Exp. Clin. Psychopharmacol. 28, 404–416. 10.1037/pha0000358

Strickland, J.C., Stoops, W.W., 2017. Stimulus selectivity of drug purchase tasks: A preliminary study evaluating alcohol and cigarette demand. Exp. Clin. Psychopharmacol. 25, 198–207. 10.1037/pha0000123

Tatara, E., Ozik, J., Pollack, H.A., Schneider, J.A., Friedman, S.R., Harawa, N.T., Boodram, B., Salisbury-Afshar, E., Hotton, A., Ouellet, L., Mackesy-Amiti, M.E., Collier, N., Macal, C.M., 2024. Agent-based model of combined community- and jail-based take-home naloxone distribution. JAMA Netw. Open 7, e2448732. 10.1001/jamanetworkopen.2024.48732

Tremblay, A., Bellisle, F., 2015. Nutrients, satiety, and control of energy intake. Appl. Physiol. Nutr. Metab. 40, 971–979. 10.1139/apnm-2014-0549

van Steenbergen, H., Eikemo, M., Leknes, S., 2019. The role of the opioid system in decision making and cognitive control: A review. Cogn. Affect. Behav. Neurosci. 19, 435–458. 10.3758/s13415-019-00710-6

Winger, G., Galuska, C.M., Hursh, S.R., 2007. Modification of ethanol’s reinforcing effectiveness in rhesus monkeys by cocaine, flunitrazepam, or gamma-hydroxybutyrate. Psychopharmacology (Berl.) 193, 587–598. 10.1007/s00213-007-0809-9

Winger, G., Galuska, C.M., Hursh, S.R., Woods, J.H., 2006. Relative reinforcing effects of cocaine, remifentanil, and their combination in rhesus monkeys. J. Pharmacol. Exp. Ther. 318, 223–229. 10.1124/jpet.105.100461

Zernig, G., Ahmed, S.H., Cardinal, R.N., Morgan, D., Acquas, E., Foltin, R.W., Vezina, P., Negus, S.S., Crespo, J.A., Stockl, P., Grubinger, P., Madlung, E., Haring, C., Kurz, M., Saria, A., 2007. Explaining the escalation of drug use in substance dependence: models and appropriate animal laboratory tests. Pharmacology 80, 65–119. 10.1159/000103923

Zhang, Y., Brownstein, A.J., Buonora, M., Niikura, K., Ho, A., Correa da Rosa, J., Kreek, M.J., Ott, J., 2015. Self administration of oxycodone alters synaptic plasticity gene expression in the hippocampus differentially in male adolescent and adult mice. Neuroscience 285, 34–46. 10.1016/j.neuroscience.2014.11.013

Zhong, X., Li, X., Mangoni, S., 2023. A Review of Agent-Based Modeling Applications in Substance Abuse Policy Research, in: 2023 Winter Simulation Conference (WSC). IEEE, pp. 150–161. 10.1109/WSC60868.2023.10407530

