## Supplement for "Quantitative Economon Model of Transactions for Drugs and Other Commodities"

**Analysis of Archival Data**

We used archival data to provide an initial examination of whether changes in demand as a function of deprivation and satiation occur due to changes in price sensitivity (i.e., demand elasticity, α) or base consumption (i.e., demand at 0 price, *Q_0_*). In one experiment, rhesus monkeys responded on progressive-ratio (PR) schedules in closed economies under different conditions of food deprivation (Hursh et al., 1989). During baseline, subjects earned all of their food from the PR schedule. In two subsequent conditions, subjects were provided with either 1/3 or 2/3 of their baseline food for ‘free’ (i.e., food was delivered independent of the PR schedule at 1800 hrs). In another condition, subjects had access to food on a fixed-ratio 1 (FR 1) schedule during 5-minute periods at the end of a PR schedule work session. In other words, after responding on the PR schedule, subjects could earn any amount of food within the 5-minute period at a “cheap” price (1 response/food pellet). Demand curves fit to these data indicated that as the amount of free/cheap food increased, elasticity of demand α systematically increased. In contrast, base consumption *Q_0_* remained relatively similar across conditions except in the 2/3 free food condition in which *Q_0_* decreased. In another experiment with heroin users, demand for hydromorphone, a mu-opioid receptor agonist, was examined under different levels of satiation (Greenwald and Hursh, 2006). Participants responded on PR schedules for hydromorphone under conditions in which they received 0 mg, 12 mg, or 24 mg of a pre-session supplemental dose of “free” hydromorphone. Demand curves fit to these data indicated that higher doses of the pre-session hydromorphone supplement were associated with systematic increases in elasticity of demand α, whereas peak consumption *Q_o_* remained relatively similar across conditions. Based on these reviewed studies, we focused here on modeling the effect of satiation and deprivation on demand as occurring through changes in demand elasticity α (i.e., Equations 4 & 5 in the main text).

**Supplemental Figures from Simulation**

**Fig. S1**

*Purchase Probabilities for Full Simulation*

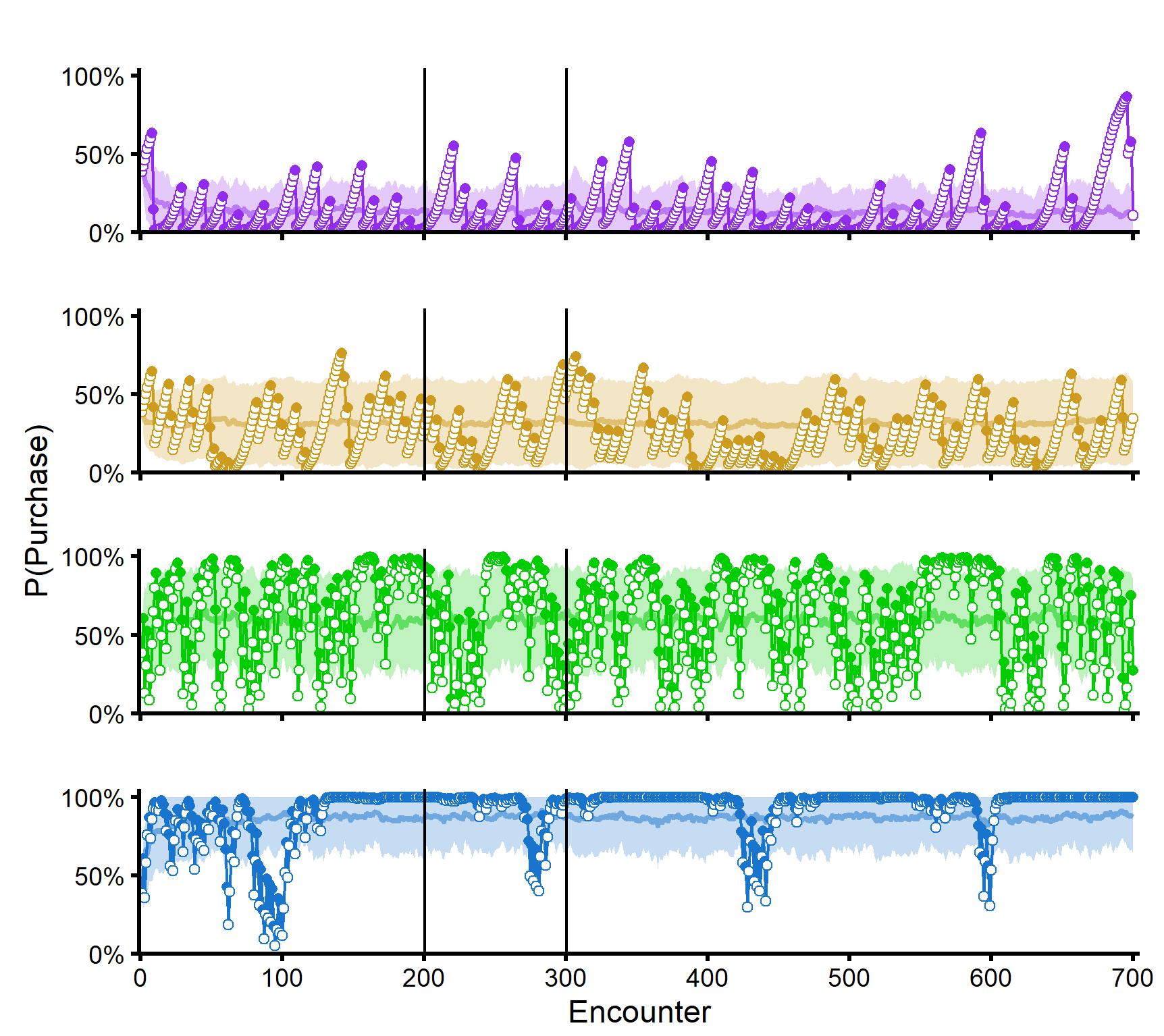

**A. Low Dep / High Sat**

**B. Low Dep / Low Sat**

**C. High Dep / High Sat**

**D. High Dep / Low Sat**

*Note.* Probability of the simulated buyers purchasing a commodity as a function of encounters with a supplier across the full simulation (700 encounters). Purchase probabilities changed according to failed and successful transactions. Failed transactions caused increases in demand for the commodity (due to deprivation); successful transactions caused decreases in demand for the commodity (due to satiation). Light datapaths represent group level (i.e., mean) purchase probabilities from 100 buyers across the 700 encounters with suppliers. Shaded areas represent ±SD around the means. Darker datapath represents purchase probabilities from one representative buyer within each condition (see main text), with open and closed symbols indicating failed and successful transactions, respectively. Transactions 1-200 were preconditioning (i.e., “burn-in”). Vertical bars indicate which subset of transaction encounters were presented in the main text (i.e., encounters 201-300).

**Fig. S2**

*Animation of Demand During Simulation*

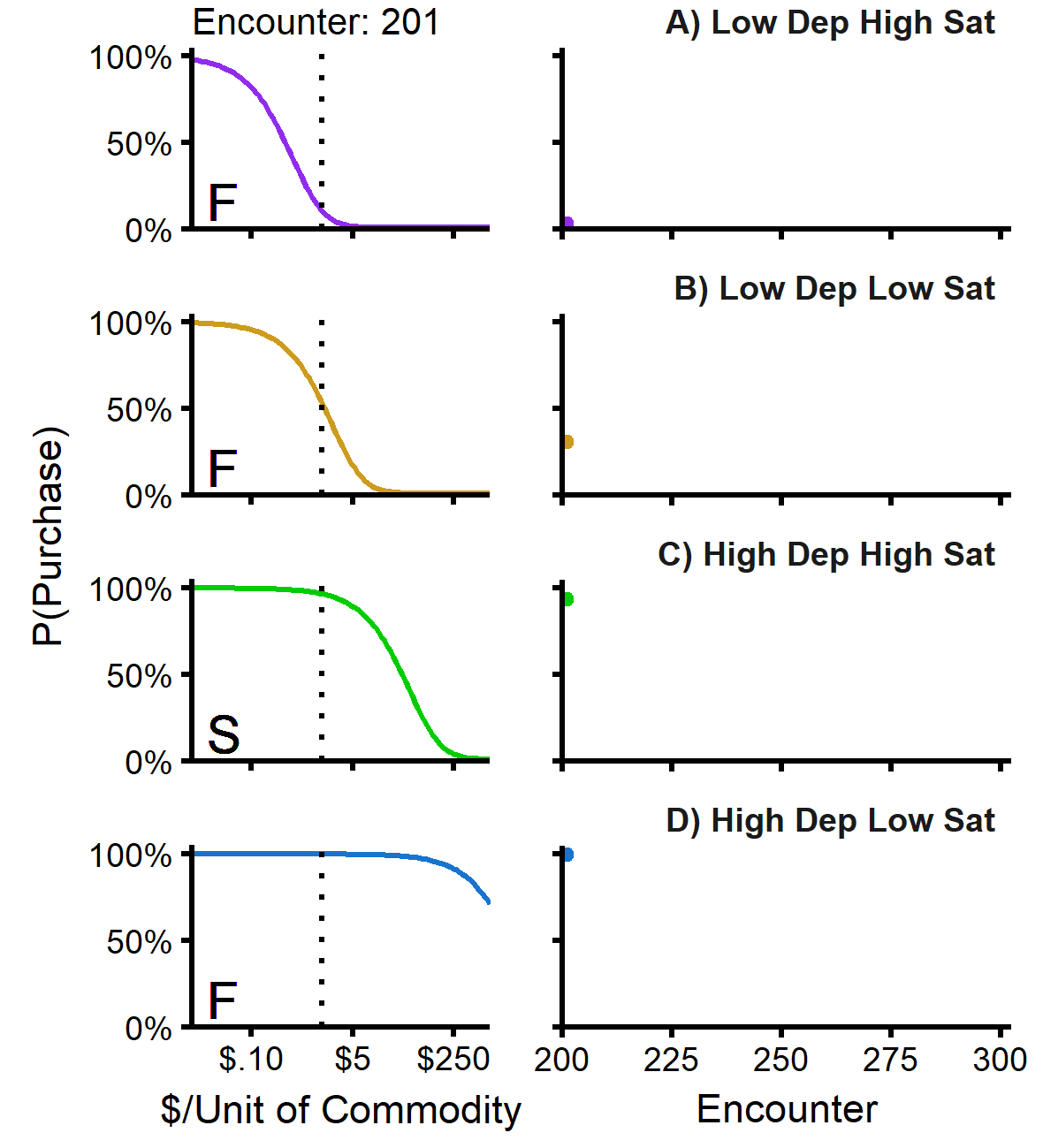

*Note.* Animation of the sequential changes in demand (i.e., probability of purchase) for the representative buyers from Fig. S1 during the encounters presented in the main text (i.e., encounters 201-300). Left: Buyer demand curves across encounters, derived from Equation 1 using *α_B,i_* and *Q_B0_* = 100%. Price of the commodity during the encounter is represented by the dashed vertical line which also represents the price at ½ *Q_0S_*. Transaction outcomes were determined at this price for all simulations. F: failed transaction; S: successful transaction. Right: Probability of the simulated buyers purchasing a commodity as a function of encounters with a supplier. Datapoint follows purchase probabilities at the price of the commodity according to the demand curves displayed in the left column.

**Table S1**

*Alternative/Future Directions for Quantitative Economon Model*

| Initial Implementation | Current Variable | Alternative Implementation | Alternative Variable |
| --- | --- | --- | --- |
| Supplier elasticity (*α_S_*) held constant. | $\alpha_{S}=.005$  $\delta_{-\alpha S}=0$  $\delta_{+\alpha S}=0$ | Transaction outcomes affect *α_S_* via satiation and/or deprivation. | If *Y_B,i_* = 1 & *Y_S,i_* = 1,  $\alpha_{S,i+1}=\alpha_{S,i}.(1+\delta_{S,+\alpha})$  else,  $\alpha_{S,i+1}=\alpha_{S,i}.(1-\delta_{S,-\alpha})$ |
| Peak purchase/sell probabilities set to 100% likelihood of purchasing/selling commodity. | $Q_{B0}=100\%$  $Q_{S0}=100\%$ | Peak purchase/sell probabilities set less than 100% likelihood of purchasing/selling commodity. | $Q_{B0}<100\%$  $Q_{S0}<100\%$ |
| Peak purchase/sell probabilities unaffected by transaction outcomes. | $Q_{B0,i+1}=100\%$  $Q_{S0,i+1}=100\%$ | Peak purchase/sell probabilities vary based on transaction outcomes. | $Q_{B0,i+1}=f(Q_{B0,i-1},Y_{B,i},Y_{S,i})$  $Q_{S0,i+1}=f(Q_{S0,i-1},Y_{B,i},Y_{S,i})$ |
| Deprivation (*δ_-ɑ_*) and satiation (*δ_+ɑ_*) parameters constant across encounters. | $\delta_{-\alpha,i+1}=\delta_{-\alpha,i}$  $\delta_{+\alpha,i+1}=\delta_{+\alpha,i}$ | Deprivation (*δ_-ɑ_*) and satiation (*δ_+ɑ_*) parameters vary based on previous transactions (e.g., inducing tolerance/dependence). | $\delta_{-\alpha,i+1}=f(\delta_{-\alpha,i+1},Y_{B,i},Y_{S,i})$  $\delta_{+\alpha,i+1}=f(\delta_{+\alpha,i+1},Y_{B,i},Y_{S,i})$ |
| Economon participants do not interact with participants outside of economon. | N/A | Economa interact as part of convergent or divergent networks. | Interactive economa: One buyer and *n* > 1 suppliers for different products (e.g., drugs vs. food) or similar products (e.g., two different drugs). |
| Deprivation and satiation used as “proof of concept” variables. | *δ_+ɑ_*  *δ_-ɑ_* | Use alternative biobehavioral or economic variables. | Examples of time-dependent variables: withdrawal severity, blood levels of drug, available income. |
| Sequence-based modeling in which demand changes based on immediately prior interactions. | If *Y_B,i_* = 1 & *Y_S,i_* = 1,  $\alpha_{i+1}=\alpha_{i}.(1+\delta_{+\alpha})$  else,  $\alpha_{i+1}=\alpha_{i}.(1-\delta_{-\alpha})$ | Time-based modeling in which demand changes based on time (*t*) elapsed since last encounter. | For deprivation,  $\alpha_{t}=\alpha_{i}.\left( 1-\delta_{-\alpha} \right)^{t}$  For satiation  $\alpha_{t}=\alpha_{i}.\left( 1+\delta_{+\alpha} \right)^{t}$ |

*Note.* Summary of future directions based on current implementation in the QEM and alternative implementation.

**Supplementary References**

Greenwald, M.K., Hursh, S.R., 2006. Behavioral economic analysis of opioid consumption in heroin-dependent individuals: effects of unit price and pre-session drug supply. Drug Alcohol Depend. 85, 35–48. https://doi.org/[10.1016/j.drugalcdep.2006.03.007](http://dx.doi.org/10.1016/j.drugalcdep.2006.03.007)

Hursh, S.R., Raslear, T.G., Bauman, R., Black, H., 1989. The quantitative analysis of economic behavior with laboratory animals, in: Understanding Economic Behaviour. Springer Netherlands, Dordrecht, pp. 393–407. https://doi.org/[10.1007/978-94-009-2470-3_22](http://dx.doi.org/10.1007/978-94-009-2470-3_22)
